# Pertpy: an end-to-end framework for perturbation analysis

**DOI:** 10.1101/2024.08.04.606516

**Authors:** Lukas Heumos, Yuge Ji, Lilly May, Tessa Green, Xinyue Zhang, Xichen Wu, Johannes Ostner, Stefan Peidli, Antonia Schumacher, Karin Hrovatin, Michaela Müller, Faye Chong, Gregor Sturm, Alejandro Tejada, Emma Dann, Mingze Dong, Mojtaba Bahrami, Ilan Gold, Sergei Rybakov, Altana Namsaraeva, Amir Moinfar, Zihe Zheng, Eljas Roellin, Isra Mekki, Chris Sander, Mohammad Lotfollahi, Herbert B. Schiller, Fabian J. Theis

## Abstract

Advances in single-cell technology have enabled the measurement of cell-resolved molecular states across a variety of cell lines and tissues under a plethora of genetic, chemical, environmental, or disease perturbations. Current methods focus on differential comparison or are specific to a particular task in a multi-condition setting with purely statistical perspectives. The quickly growing number, size, and complexity of such studies requires a scalable analysis framework that takes existing biological context into account. Here, we present pertpy, a Python-based modular framework for the analysis of large-scale perturbation single-cell experiments. Pertpy provides access to harmonized perturbation datasets and metadata databases along with numerous fast and user-friendly implementations of both established and novel methods such as automatic metadata annotation or perturbation distances to efficiently analyze perturbation data. As part of the scverse ecosystem, pertpy interoperates with existing libraries for the analysis of single-cell data and is designed to be easily extended.

## Introduction

Understanding cellular response to stimuli is crucial for identifying biological phenomena and mechanisms. Single-cell data has increasingly shifted from observational experiments to perturbation experiments, encompassing genetic modifications, chemical treatments, physical interventions, environmental changes, diseases, and combinations thereof. Technologies such as Perturb-seq^1^, CROP-seq^2^, and Sci-plex^3^ leverage single-cell readouts to capture perturbations at scale. By monitoring resulting shifts in intrinsic cell states, single-cell perturbation analyses offer insights into changes in gene programs, shared and divergent responses across tissues, drug targets and interactions, changes in cell type frequency, and in cell-cell interactions after perturbation.

Statistical and machine-learning based analysis methods have been developed for such complex data^4–7^ resulting in the discovery of, for example, cell states associated with autism risk genes^8^ or stimulation responses in primary human T cells^9^. However, the size and complexity of larger perturbation screens can pose significant challenges for interpretation without meaningful lower-dimensional representations and additional context regarding cell lines or perturbations. No current analysis framework exists which scales to genome-scale datasets^10^, contextualizes data with public annotations, and uses common data structures across tools. Furthermore, many highly single-task tools suffer from maintenance issues, or are confined to the R ecosystem, complicating the analysis. We have seen from other, widely used frameworks within the single-cell realm, such as scirpy^11^ for adaptive immune receptor data or scvi-tools^12^ for probabilistic modeling, to enable the efficient analysis of multimodal data while providing building blocks for developers to build upon. Inspired by their impact and the lack of efficient frameworks for perturbation data, we develop a new framework focused on perturbation data within scverse^13^.

Pertpy, a framework for **pert**urbation analysis in **Py**thon, is purpose-built to organize, analyze, and visualize complex perturbation datasets. Pertpy is flexible, and can be applied to datasets of different assays, data types, sizes, and perturbations, thereby unifying previous data-type or assay-specific single-problem approaches. Designed to integrate external metadata with measured data, it enables unprecedented contextualization of results through swiftly built, experiment-specific pipelines, leading to more robust outcomes. To evaluate methods and obtained representations for perturbations, we implemented a series of shared metrics. The wide array of use-cases and different types of growing datasets are addressed by pertpy through its sparse and memory-efficient implementations, which leverage the parallelization and GPU acceleration library Jax^14^, thereby making them up to eight times faster than original implementations (**Supplementary Figure 1**). We demonstrate this versatility by applying pertpy to three different, popular scRNA-seq perturbation use-cases. To show how pertpy can discover new gene programs, we remove confounding factors in a CRISPR screen (Perturb-seq)^15^ study to project it into a meaningful perturbation space, where we transfer labels of known to unknown gene programs. Moreover, we demonstrate how pertpy can be used to deconvolve perturbation responses into viability-dependent and -independent components in a large-scale gene expression and drug response screen^16^ by integrating metadata from existing databases. Finally, we decipher compositional changes, rank perturbation effects, and find treatment response-specific multicellular programs in a triple negative breast cancer study^17^. Whereas previously a user would separately download cell line or perturbation information from scattered databases while piecing together analysis tools from different, incompatible ecosystems, it is now possible to efficiently analyze complex perturbation datasets end-to-end with integrated biological context.

We provide online links to tutorials with more than 15 additional use-cases that demonstrate pertpy’s usage with datasets spanning a variety of cell lines and perturbation conditions, ranging from CRISPR screens^18^ and exposure to pathogens^19^ to inflammation^20^ and COVID-19 severity states^21^. Pertpy is accessible as an extendable, user-friendly open-source software package hosted at https://github.com/scverse/pertpy and installable from PyPI and Conda. It comes with comprehensive documentation, tutorials and use-cases available at https://pertpy.readthedocs.io.

## Results

### Pertpy enables fast and scalable perturbation analysis

Pertpy includes methods for single and combinatorial perturbations to cover diverse types of perturbation data including genetic knockouts, drug screens, and disease analyses. The framework is designed for flexibility, offering more than 100 composable and interoperable analysis functions organized in modules which further eases downstream interpretation and visualization. These modules host fundamental building blocks for the implementation and methods that share functionality that can then be chained into custom pipelines. To facilitate setting up these pipelines, pertpy guides analysts through a general analysis pipeline (**Figure 1**) with the goal of elucidating underlying biological mechanisms by examining how specific interventions alter cellular states and interactions. In a data transformation step, any confounding factors and artifacts are first removed during quality control. To identify and visualize groups of similarly behaving perturbations, a perturbation space, which quantifies data similarities grouped by phenotype, can be calculated that can more easily be interpreted by enriching the cell lines or perturbations with metadata. In a second knowledge inference step, the obtained representations are used as input for statistical and machine learning models.

**Figure 1.**
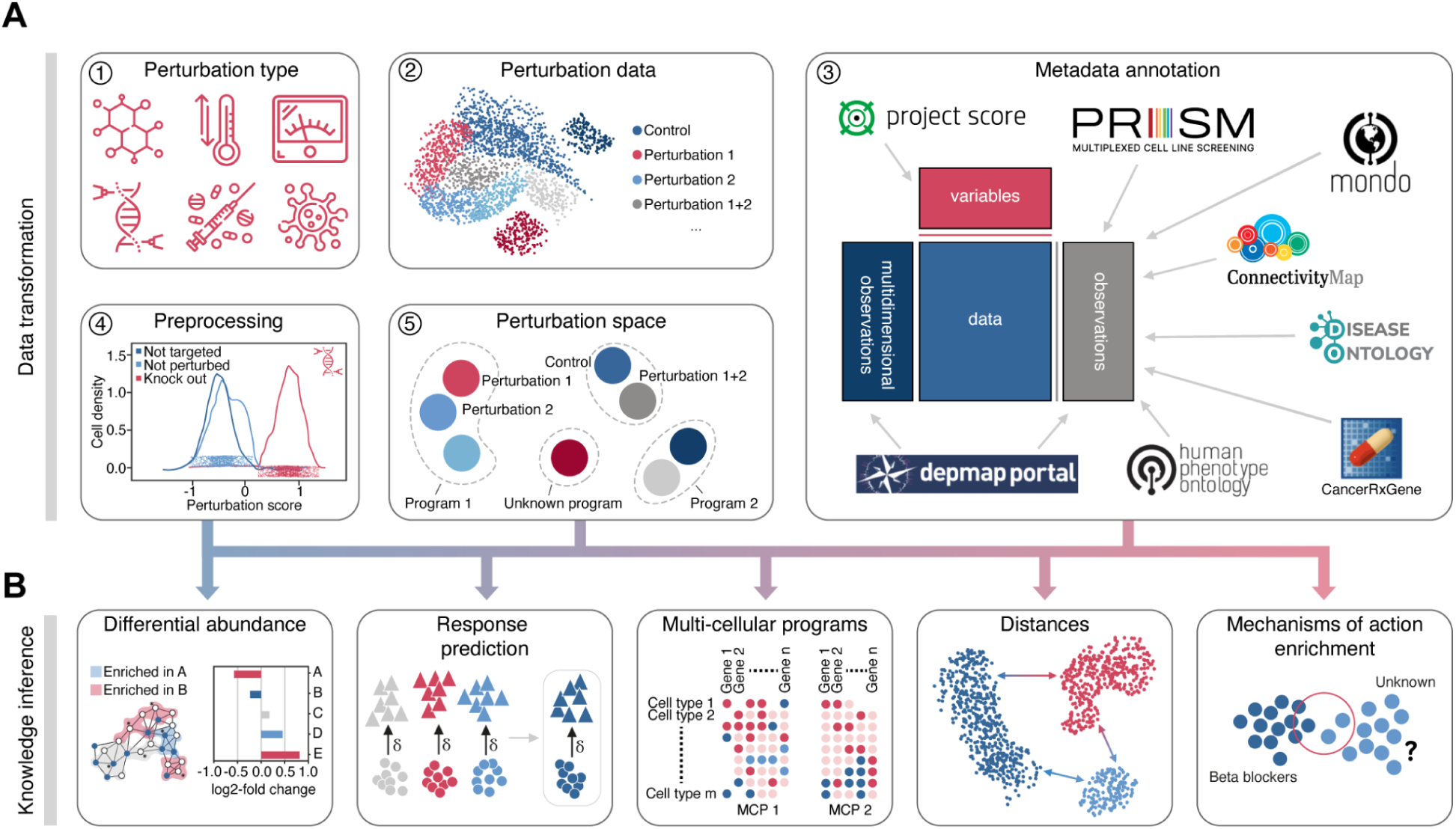
Modules of the pertpy framework. (A) Unimodal or multimodal single-cell perturbation data originating from genetic modifications, chemical treatments, physical interventions, environmental changes, or diseases is enriched with metadata from several databases. During preprocessing, confounding factors such as cell cycle and batch effects may be removed. Targeted cells are labeled as successfully or not successfully perturbed. Altogether this enables the calculation of a meaningful perturbation space. (B) Pertpy enables downstream analyses, dependent on the question of interest. These include differential analysis, response prediction, the determination of multicellular programs, the calculation of distance between perturbations, and mechanism of action enrichment.

The input to a typical analysis with pertpy are unimodal scRNA-seq or multimodal perturbation readouts stored in AnnData^22^ or MuData^23^ objects. While pertpy is primarily designed to explore perturbations such as genetic modifications, drug treatments, exposure to pathogens, and other environmental conditions, its utility extends to various other perturbation settings, including diverse disease states where experimental perturbations have not been applied. A typical analysis with pertpy starts by curating the perturbation metadata against ontologies such as the Cell Line ontology^24^ or the Drug ontology^25^ and annotating the perturbations with additional metadata obtained from Depmap and Genomics of Drug Sensitivity in Cancer (GDSC)^26^ for cell lines, the Connectivity Map (CMap)^27^ for mechanisms of action, and the pubchem^28^ and CHEMbl^29^ databases for drugs (**Methods**). Next, if the data originates from a CRISPR screen, pertpy assigns guide RNAs to cells. The application of CRISPR can exhibit variable efficacy in affecting gene expression. Pertpy’s fast mixscape^18^ implementation accounts for this by classifying targeted cells based on their response to a perturbation, analyzing each cell’s perturbation signature to determine if they were successfully perturbed (**Methods, Supplementary Figure 1**). As the number of applied perturbations increases, it becomes increasingly challenging to compare and interpret them further. Pertpy provides several distinct ways to learn biologically interpretable perturbation spaces that depart from the individualistic perspective of cells and instead generate a single embedding per perturbation to summarize the cellular responses (**Methods**). This specialized space enables representing the collective impact of perturbations on cells and serves as potential input for downstream analysis methods^15,30,31^. Generally, pertpy’s analysis pipeline can be adapted depending on whether the experiment involved multiple cell types or a number of experimental perturbations.

To robustly identify biological variation across conditions, pertpy fills a gap within the single cell analysis ecosystem by providing an intuitive interface for differential gene expression that supports complex designs and contrasts which is needed for multi-condition data and natively supported in scanpy (**Methods**). Currently, pertpy supports PyDESeq2^32^, edgeR^33^, Wilcoxon, and T-tests. Going beyond differential gene expression at scale, both annotated metadata and differentially expressed genes can then be used as input for further pertpy modules such as gene set enrichment tests to uncover the biological effects induced by the perturbations (**Methods**).

Tracking cell type compositional shifts is crucial for understanding the underlying mechanisms of disease progression, tissue regeneration, and developmental biology, offering insights into cellular responses and adaptations. Pertpy offers two distinct approaches for detecting compositional shifts, both utilizing a common MuData-based data structure. If labeled groups are available, pertpy provides substantially accelerated and more scalable implementations of scCODA^34^ 2.0 and its cell type hierarchy-aware extension tascCODA^35^ 2.0 (**Supplementary Figure 1**, **Methods**). Both methods employ Bayesian methods to elucidate cell type compositional changes. If no labeled groups are available or continuous proportions are expected, such as during developmental processes, pertpy implements a scalable version of Milo, originally unique to the R ecosystem^36^, to conduct differential abundance tests by assigning cells to overlapping neighborhoods within a k-nearest neighbor graph (**Methods**).

Understanding how cells function together within tissues is a significant challenge. Multicellular programs (MCPs) refer to the orchestrated activities of various cell types that collaborate to create complex functional structures at the tissue scale. Pertpy’s fast and scalable implementation of DIALOGUE^37^ uncovers MCPs through a combination of factor analysis and hierarchical modeling, thanks to a novel fast input-order invariant linear programming solver and a new, fast test to determine significantly associated MCP genes (**Methods**).

Not all cell types are equally affected by perturbations. Pertpy’s fast implementation of Augur (**Supplementary Figure 1)** ranks cell types based on their response to perturbations by training machine learning models to predict experimental labels within each cell type, and then ranking these cell types by the models’ accuracy metrics across multiple cross-validation runs (**Methods**). Further, understanding the dynamics of cellular responses to various stimuli is crucial in particular when the experimental exploration of all possible conditions is unfeasible.

CINEMA-OT^38^, as implemented in a scalable manner in pertpy, extends this concept by distinguishing between confounding variations and the effect of perturbations, achieving an optimal transport match that mirrors counterfactual cell pairings (**Methods**). These pairings enable analysis of potentially causal perturbation responses, allowing for individual treatment-effect analysis, clustering of responses, attribution analysis, and the examination of synergistic effects.

For accurate statistical comparison and measurement of perturbation effects, it is essential to employ distance metrics between cell groups. A suitable metric quantifies divergence or similarity in expression patterns of cells under different perturbations, allowing to infer unique or common mechanisms. Different types of distance metrics make varying assumptions on the shape of the data and emphasize specific aspects of it. For instance, optimal transport based distances such as the Wasserstein distance assume correspondence between cell populations, while Mahalanobis distance focuses on covariance structures and scale differences within the data. In order to capture a wide range of distance metric types, more than 18 different metrics including, but not limited to, the euclidean distance, E-distance^39^, and the Wasserstein distance^40^ are implemented in pertpy (**Methods**). All included metrics can also be used for perturbation testing through Monte-Carlo permutation testing, allowing for the statistical evaluation of perturbation distinguishability and efficacy (**Methods**).

Building upon the scverse^13^ ecosystem, pertpy ensures interoperability with existing workflows for single-cell omics analyses and can be purposefully extended to solve new challenges. Base classes for additional perturbation spaces, distances, differential gene expression tests, and other components are provided to facilitate swift development. We additionally provide a dataset module with more than 30 public loadable perturbational single-cell datasets in MuData and AnnData format building upon and extending scperturb^39^ to kickstart analysis, development and benchmarking with pertpy. The metadata of the datasets were curated against public ontologies to enable swift dataset integration and large-scale machine learning including foundational models.

### Pertpy facilitates the exploration of genetic interaction manifolds

To demonstrate pertpy’s ability to create meaningful perturbation spaces, we examined a publicly available CRISPR screen dataset initially presented by Norman et al.^15^, consisting of 111,255 single-cell transcriptomes of K562 cells subjected to 287 single gene and gene pair perturbations. This dataset allows us to investigate how genetic interactions through combinatorial expression of genes lead to cellular and organismal gene programs and phenotypes.

After preprocessing (**Methods**), we applied pertpy’s mixscape^18^ implementation to remove confounding effects such as the cell-cycle by calculating cell-specific perturbation signatures (**Figure 2A, Methods**). To identify targeted cells that escaped gene knockout despite the presence of a guide RNA and should thus not be considered perturbed, we applied mixscape’s mixed effect model, which improved the Silhouette Coefficient for the perturbation label by 0.097. Specifically, we identified 25,183 such cells **(Figure 2B),** characterized by a low perturbation score that we removed in contrast to the original authors, compared to 74,237 cells with sufficient signal **(Figure 2C)** which we kept for further analysis.

**Figure 2.**
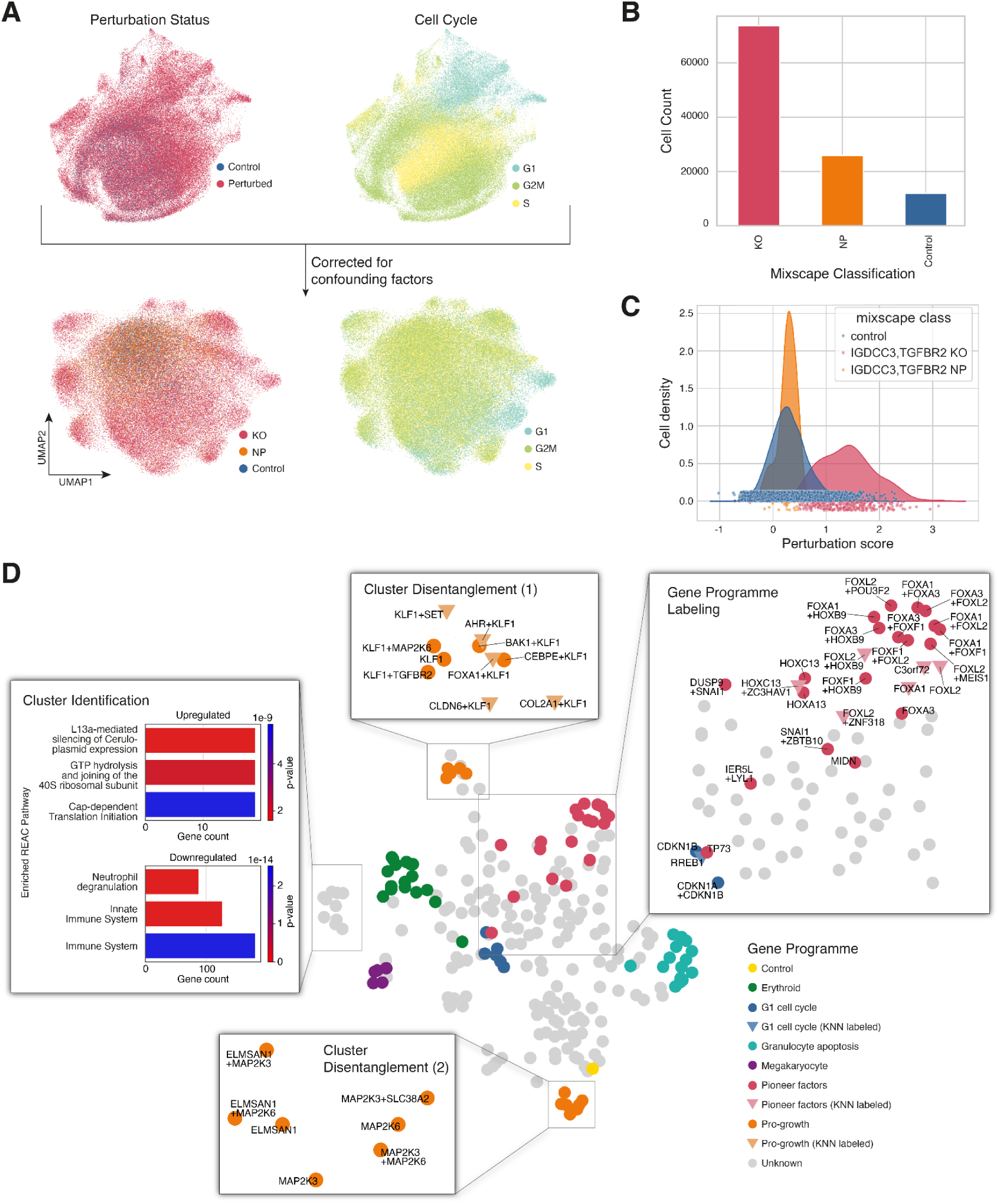
Learning a unified perturbation space in combinatorial CRISPR perturbation scRNAseq data^15^ via pertpy’s perturbation space pipeline. Before preprocessing, the dataset featured control, targeted but not successfully perturbed (NP), and successfully perturbed (KO) cells. (A) Pertpy’s mixscape implementation removes confounding factors such as cell cycle effect. (B) Pertpy’s application of mixscape determined 25,183 cells to be targeted but labeled as not successfully perturbed. (C) Example density plot of a combination gene knockout. (D) Perturbation space highlighting gene programmes that were originally labeled by Norman et al.^15^ (solid colors, circles) and determined via label transfer (opaque colors, triangles).

We then projected the perturbation signature into a perturbation space using the penultimate layer of our multi-layer perceptron based discriminator classifier (**Figure 2D, Methods**). Importantly, we observe that explicitly training the classifier to distinguish between individual perturbations results in the clustering of perturbations with similar effects on the cell, as indicated by the affected gene programme as originally labeled by Norman et al.^15^. In addition to validating known annotations, pertpy also enables extending these to clusters with unknown gene programmes using k-nearest neighbor based label transfer (**Methods**). Evaluating data in perturbation space also allows for a refinement of previous annotations. For instance, the perturbation TP73, characterized as a pioneer factor gene programme in the original publication^15^, clusters with the G1 cell cycle perturbations when embedded using the discriminator classifier. This can be explained by TP73’s profound influence on the cell cycle^41^. Moreover, what the original authors identified and labeled as a single pro-growth gene programme cluster can now be differentiated into two distinct clusters. Indeed, we found that although both clusters comprise perturbations targeting genes important for cell growth, one cluster mainly targets Krüppel-like factors (KLFs), while the other cluster comprises cells with perturbed mitogen-activated protein kinases (MAPK). In summary, the projection of data into the perturbation space also allows for an in-depth exploration of clusters without gene programme annotation, enabling the identification of a novel, previously unannotated cluster, which comprises perturbations with a profound downregulating effect on the neutrophil degranulation pathway (**Figure 2D**). This use-case demonstrates the simplicity and effectiveness of combining several of pertpy’s modules into a new analysis pipeline from quality control over perturbation space to the annotation of previously unlabeled gene programs.

### Pertpy streamlines discovery for complex perturbation experiments

Advancements in multiplexing technologies have significantly increased the number of cell states which can be profiled in one experiment, resulting in large perturbation screens. McFarland et al.^16^ introduced MIX-Seq, an experimental assay which enables the multiplexing of different cell lines within a single sequencing run^16^. We use pertpy to efficiently analyze a dataset comprising 172 cell lines and 13 drug treatments^16^.

Pertpy reduces annotation and quality control to just a few steps. Its metadata module annotates the cell lines with tissue-of-origin, cancer type, and bulk expression profiles from the disease ontology Oncotree^42^ and the Cancer Cell Line Encyclopedia^43^ (CCLE). Compounds are annotated with compound targets and mechanism of action from the Cancer Dependency Map (DepMap)^44^, Genomics of Drug Sensitivity in Cancer^26^ (GDSC), and Connectivity Map (CMap)^27^ (**Methods**). Following annotation, pertpy enables immediate visualization for exploratory analysis (**Figure 3A**). Additionally, annotated bulk expression allows users to compare RNA profiles of their cell lines with established public datasets, providing rapid quality control functionality. Comparative analysis revealed an average Pearson correlation coefficient of 0.88 across all cell lines (**Figure 3B**), demonstrating substantial consistency with the cell line passages cataloged in the DepMap CCLE database, and enabling the integration of additional screening data from the DepMap PRISM project^31^.

**Figure 3.**
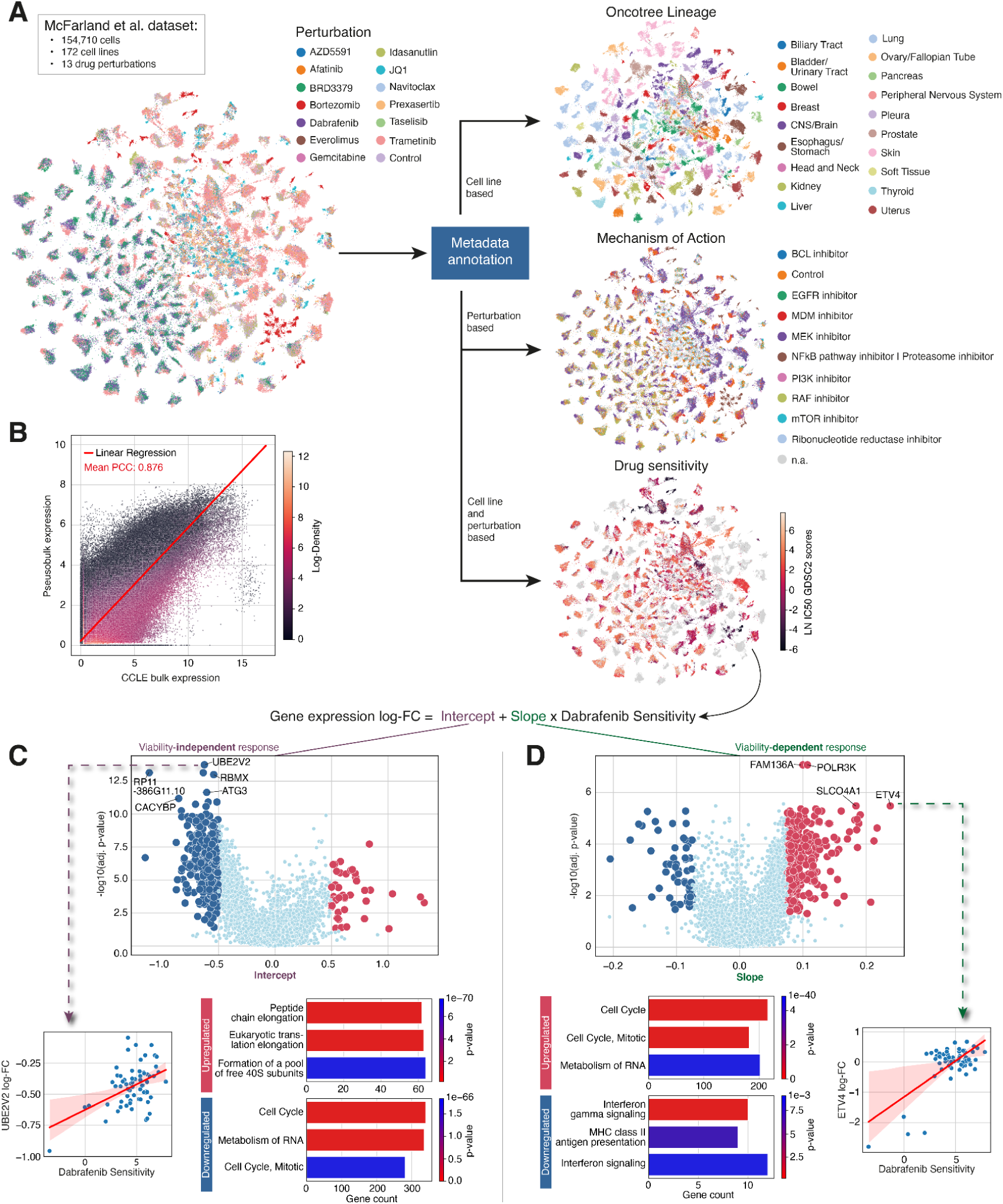
Deconvolution of viability-related response signatures in scRNA-seq drug screen data^16^. (A) Overview of the chemical perturbation dataset. Cell lines and perturbations were annotated with pertpy with additional metadata facilitating detailed analysis. (B) Linear regression model between single-cell expression data and GDSC profiles show high correlation reinforcing the quality of the dataset. (C) Volcano plot showing the value and significance (Benjamini-Hochberg corrected) of the intercept of the fit linear regression models for each gene (top), indicating the viability-independent response. An example linear regression for the gene UBE2V2 (bottom left) shows that a change in UBE2V2 expression in a cell line is observable, irrespective of the respective cell line’s sensitivity to dabrafenib treatment. The top genes were used to perform gene set enrichment analysis (bottom right). The figure design is inspired by Figure 2C in the original paper^16^. (D) Same as (C), but for the slope of the linear regression models, indicating the viability-dependent response.

Pertpy significantly streamlines the replication and extension of the original analyses by McFarland et al.^16^. We use pertpy to fetch and annotate IC50 values for each cell line and perturbation pair from GDSC (**Methods**). This allows us to easily replicate the original statistical method to uncover viability-dependent and -independent gene expression associations. We selected a different drug from the original analysis^16^, the BRAF inhibitor dabrafenib^45^, and used pertpy to compute post-treatment log-fold changes across 81 cell lines (**Methods**). We interpret the intercept and slope of the linear regression on IC50 to be the viability-independent and -dependent responses of the respective gene to dabrafenib (**Methods**, **Figure 3 C-D**). Notably, we find that genes like UBE2V2, RP11-386G11.10, and ETV4, which are linked to cancer progression^46–48^, displayed significant variations in their fitted response parameters (**Figure 3C-D**). Additionally, our analysis identified an enrichment of genes involved in interferon signaling and MHC class II antigen presentation in the viability-dependent genes, consistent with the initiation of an immune-mediated cell death response to dabrafenib (**Figure 3D**). Interestingly, protein translation pathway genes were upregulated in the viability-independent effects of dabrafenib, a response previously noted with dabrafenib^49^ but with no mechanistic information until now. This mechanism is distinctly different from dabrafenib’s putative mechanism of action, BRAF inhibition, which targets an orthogonal cell survival pathway. Pertpy’s ability to efficiently manage, analyze, and supplement complex experimental design with additional datasets underscores its utility in conducting sophisticated biology-informed analyses. This streamlined approach greatly enhances the depth of biological insights.

### Pertpy enables deciphering effects of perturbations on cellular systems

Understanding the complex interplay between the immune system and the tumor microenvironment (TME) is crucial for unraveling cancer progression. This is particularly important in solid tumor entities, such as triple-negative breast cancer (TNBC), a rare, aggressive breast cancer subtype that lacks estrogen, progesterone, and human epidermal receptors, rendering it unresponsive to standard receptor-targeted therapies^50^. Single-cell transcriptomics of breast cancer tumors has uncovered distinct T-cell subtypes and the involvement of plasmacytoid dendritic cells (pDCs) in promoting immunosuppression within the TME in TNBC through tumor-immune crosstalk^51^ which is a significant driver of treatment resistance^52^. Studies have further elucidated TNBC-specific features and differential responses to neo-adjuvant chemotherapy (NACT) and immunotherapy, highlighting the role of PD-1 and PD-L1 pathways in modulating treatment outcomes^53^. Therefore, we set out to demonstrate how pertpy can be used to investigate treatment responses using a publicly available dataset of 22 TNBC patients treated with NACT with and without additional PD-L1 inhibitor paclitaxel^17^ administration, initially presented by Zhang et al.^17^ (**Methods**, **Figure 4A-B**).

**Figure 4.**
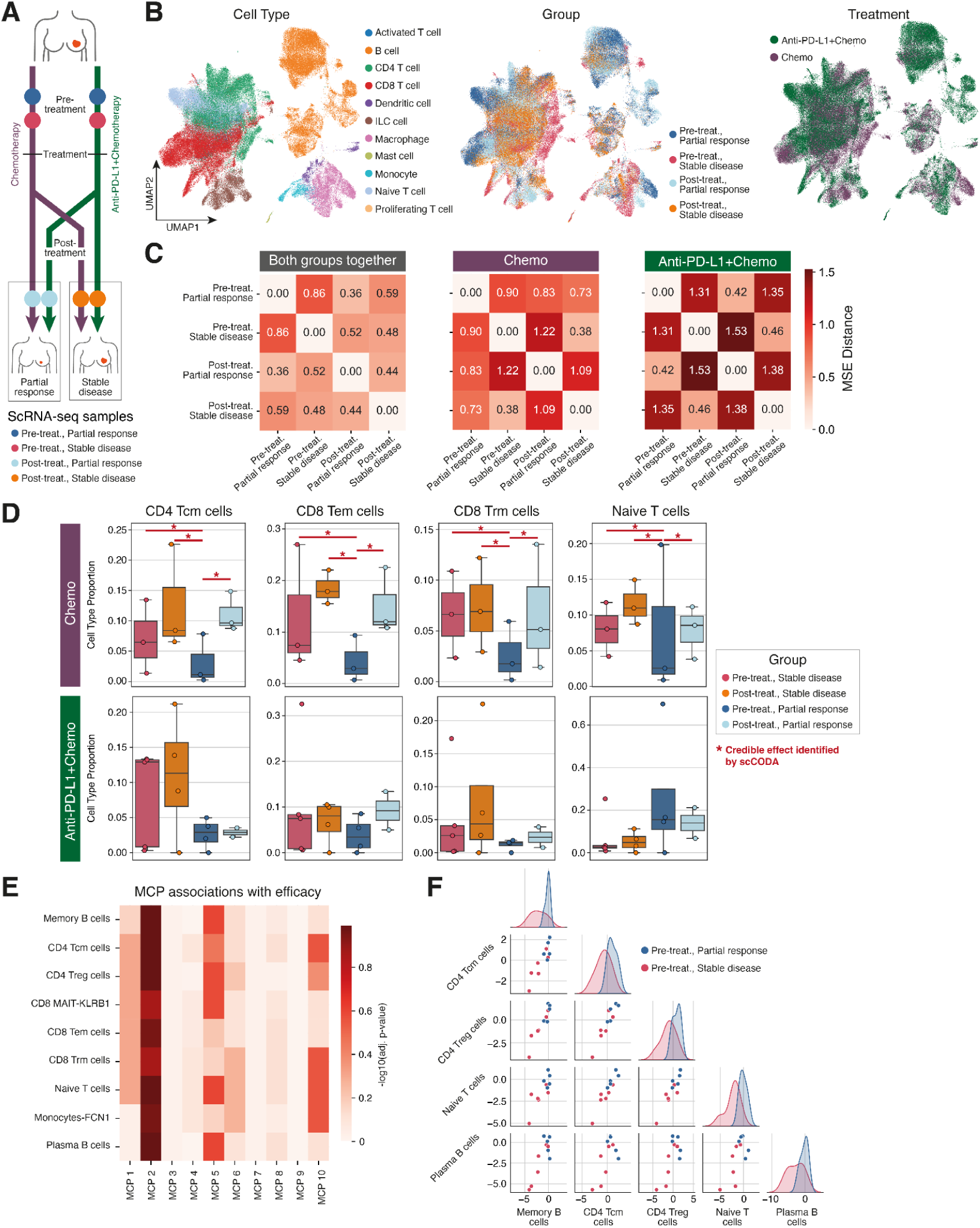
Pertpy identifies complex perturbation effects in multicellular tissue as demonstrated on a TNBC treatment dataset^17^. (A) Schematic overview of the experimental design. (B) Single-cell RNA-seq of tissue from 22 patients with triple-negative breast cancer, comparing pre- and post-treatment responses to anti-PD-L1 therapy and neoadjuvant chemotherapy. (C) Mean-squared error (MSE) distance between treatment responses shows higher distances between partial responses and stable disease. (D) scCODA analysis shows significant compositional changes for patients treated with chemotherapy. (E) DIALOGUE analysis shows several multicellular programs (MCPs) associated with treatment efficacy. (F) Pairplot of MCP 2.The diagonal shows a cell type specific kernel density estimate of the mean score for each MCP by sample. In the lower triangle’s scatter plots, each point denotes an average patient score for the cell types labeled on the corresponding row (x-axis) and column (y-axis).

To rank perturbation effects, we used pertpy to calculate the Euclidean distance between the pre- and post-treatment patients of the four groups due to its superior performance in independent benchmarks^54^. We found that patients responding to NACT alone had a greater distance between pre- and post-treatment expression profiles compared to responders to anti-PDL-1 and NACT combination therapy, implying that the latter led to potentially a less intense response or was used in cases with a worse prognosis. To identify cell types involved in treatment response, we investigated shifts in cell type composition induced by the treatment.

Tracking cell type shifts is essential for understanding disease progression, tissue regeneration, and treatment responses, revealing key insights into cellular adaptations. We applied pertpy’s fast implementation of the Bayesian model scCODA^34^ 2.0 to the dataset per treatment (**Methods**). We found compositional shifts for NACT treatment in CD4 central memory, CD8 effector memory, CD8 tissue-resident memory, and naive T cells between disease stages but not for combination therapy (**Figure 4D**). To better understand whether the cell types that are subject to compositional shifts are a part of a common cell circuit, we set out to find shared gene expression signatures in several cell types which jointly act as tissue level units, so-called multicellular programs (MCPs)^37^.

We applied pertpy’s implementation of DIALOGUE^37^, which finds MCPs using matrix decomposition in conjunction with a novel, fast input-order invariant linear programming solver, to the TNBC treatment dataset to uncover 10 multicell type signatures predictive of treatment response (**Methods**). We find that a patient’s average MCP2 score in seven different cell types (**Supplementary Table 1**) is predictive of treatment response for both treatments (adj. P≤ e-01) (**Figure 4D, Supplementary Figure 2A,B**). We pooled patients receiving both treatments for this analysis due to scalability limitations of the DIALOGUE method, which requires all cell types analyzed be present in all patients. Several of the MCP2-associated genes (**Methods, Supplementary Figure 2**) are associated with heat shock proteins (HSPs), implying a role for these proteins in immune cells in treatment response. For instance, HSPA1B, which is significantly increased in MCP2 for all tested cell types (**Methods**), has been previously identified as a prognostic biomarker in breast cancer^55,56^. Variations in HSPs within the cellular tumor microenvironment and cancer cells, potentially influenced by external factors or communication among cell types, may impact tumor progression. Therefore, we directly investigated cell-cell communication using a public ligand-receptor gene database (**Methods**). We found several interactions, such as the experimentally validated interaction between the cytokine ligand IL-7, which has a known antitumor role across diverse cancers^57^, and its receptor IL7R. The gene encoding IL-7 is a MCP 2 associated gene (**Methods**) in memory B cells and its receptor is expressed by central memory T cells and naive T cells. Increased IL-7 activity may contribute to suboptimal treatment outcomes by affecting T cell behavior and elevating levels of JUN, FOS, and FOSB, which are key components of the AP-1 complex that can either inhibit or promote tumor growth, depending on the context^58,59^. Surprisingly, lower AP-1 activity is linked to T cell exhaustion^60^, which is counterintuitive since treatment responders often exhibit greater T cell exhaustion. Whether IL-7 signaling directly causes these observations, or if a common underlying signal is influencing adaptive immune responses and IL-7 signaling requires further experimental validation.

## Discussion

Pertpy facilitates the end-to-end analysis of complex perturbation datasets with a versatile toolbox of interoperable components, encompassing metadata annotation, data analysis, and visualization tools. Through shared infrastructure and modules, we developed improved versions of widely used methods that were originally unmaintained or only easily available to the R community together with the original authors making them widely available to the community. Our community effort will ensure that all of these methods are jointly maintained and further developed. We demonstrated pertpy’s flexibility through several use-cases including the identification of perturbation-specific gene programmes using a CRISPR screen (Perturb-seq) dataset, deconvolution of viability-related response signatures in a chemical perturbation dataset, and deciphering treatment response to drugs in TBNC. Many further use-cases can be found in pertpy’s extensive tutorials.

As perturbation datasets grow larger and incorporate additional modalities like spatial transcriptomics, we anticipate the development of specialized methods for analyzing multimodal perturbation data. By combining efforts such as Squidpy^61^ and pertpy, additional functionality designed for spatial perturbations to uncover, for example, differentially regulated neighborhoods could be made widely available.

Finally, we expect pertpy to support the creation of perturbation atlases through harmonized data collection, the generation of meaningful perturbation spaces, and the evaluation of these spaces using pertpy’s distance metrics. Such atlases can comprehensively characterize cell types under various conditions to capture the wide array of inducible cell states beyond their basal states. We expect such atlases to become essential for the development of robust and generative foundation models where perturbation analysis is a key task that can be confidently evaluated with pertpy’s metrics.

We expect pertpy to lead to more robust biological discoveries through its capability of enriching measurements with biological metadata. As an extendable and interoperable framework, we anticipate pertpy to become an enabler for further robust perturbation analysis methods that tackle the growing complexity and multimodality of perturbation data.

## Methods

### Implementation of pertpy

Pertpy is implemented in Python and builds upon several scientific open-source libraries including NumPy^62^, Scipy^63^, jax^14^, scikit-learn^64^, Pandas^64,65^, AnnData^22^, scanpy^66^, muon^23^, NumPyro^67^, ott-jax^68^, blitzgsea^69^, PyTorch^70^, and scvi-tools^12^ for omics data handling, matplotlib^71^ and seaborn^72^ for data visualization.

#### Guide RNA assignment

Assigning relevant guides to each cell is essential in genetic perturbation assays, ensuring that the observed cellular responses are accurately linked to the intended genetic modifications. This step is critical for validating experimental design and interpreting results reliably. Pertpy’s module assigns cells to the most expressed guide RNA if it additionally exceeds an optional user specified count threshold.

#### Differential gene expression

Differential gene expression analysis compares the mean gene expression levels between different conditions or groups to identify genes with statistically significant changes utilizing statistical models to account for between-sample variability and control for false-discovery rates. Pertpy provides a unified application programming interface (API) to support a variety of such models. The first group of models comprises the T-test and Wilcoxon test as simple statistical tests for comparing expression values between two groups without covariates. The second group includes models of the linear model family that allow modeling complex designs and contrasts. Currently included are PyDESeq2^32^, edgeR^33^ as well as a wrapper around statsmodels (https://www.statsmodels.org) which provides access to a wide range of regression models, including ordinary least squares regression, robust linear models and generalized linear models. Linear model designs can be specified via Wilkinson formulas as known from R (through formulaic, https://github.com/matthewwardrop/formulaic). Pseudobulk workflows that account for pseudoreplication bias^73^ are enabled by integration with scanpy’s *get.aggregate()* function. Results tables ranked by adjusted p-value are provided as a pandas data frame and can be visualized using volcano plots.

#### Analysis of pooled CRISPR screens with mixscape

CRISPR-Cas9 can sometimes lead to cells escaping gene knockout by receiving an ineffective in-frame mutation, underscoring the necessity for computational quality control to predict and enhance their specificity and performance. Mixscape classifies targeted cells, i.e. those identified as perturbed by presence of a guide RNA, into successfully perturbed (KO) and targeted but not successfully perturbed (NP) based on their response. In particular, the mixscape pipeline includes the following steps:

1. Calculate the perturbation-specific signature of every cell, which is the difference of the targeted and the closest *k* (defaults to 20) nearest control neighbors.
2. Identify and remove cells that have ‘escaped’ CRISPR perturbation by estimating the distributions of KO cells. Afterwards, the posterior probability that a cell belongs to the KO cells is calculated and the cells are binary assigned based on a fixed probability threshold (defaults to 0.5).
3. Visualize similarities and differences across different perturbations using linear discriminant analysis.

We implemented mixscape following the implementation of the original authors^18^. We further optimized the implementation by using PyNNDescent (https://github.com/lmcinnes/pynndescent) for nearest neighbor search for the calculation of the perturbation signature.

#### Compositional analysis of labeled groups with scCODA and tascCODA

Tracking cell type shifts is crucial for understanding the underlying mechanisms of disease progression, tissue regeneration, and developmental biology, offering insights into cellular responses and adaptations. Despite their critical role in biological processes like disease, development, aging, and immunity, detecting shifts in cell type compositions through scRNA-seq is challenging. Statistical analyses must navigate various technical and methodological constraints, including limited experimental replicates and compositional sum-to-one constraints^34^. scCODA and its extension tascCODA both employ Bayesian methods to elucidate cell type compositional changes with tascCODA being able to also take cell type hierarchies into account.

The implementations of scCODA 2.0 and tascCODA 2.0 are mathematically equivalent to the original implementations^34,35^, but allow for accelerated inference by replacing the Hamiltonian Monte Carlo algorithm in TensorFlow^74^ with the no-U-turn sampler from numpyro^67^. The joint implementation also allows users to conveniently apply both methods from within the same framework.

Pertpy further uses MuData^23^ objects to simultaneously handle cell-by-gene and sample-by-cell type representations of the same data, simplifying the data aggregation and model specification steps for scCODA 2.0 and tascCODA 2.0 while ensuring compatibility with other methods featured in the scverse^13^ ecosystem. A wide range of visualization options through scanpy^66^, ete3^75^, and arviz^76^ for representation of differentially abundant cell types, their hierarchical structure, and inference diagnostics respectively are also provided within pertpy.

#### Compositional analysis of unlabeled groups with Milo

Most methods for comparing single-cell datasets often rely on identifying discrete clusters to test for differences in cell abundance across experimental conditions. Yet, this approach may lack the necessary resolution and fail to represent continuous biological processes accurately. To address these limitations, Milo was designed to conduct differential abundance tests by assigning cells to overlapping neighborhoods within a k-nearest neighbor graph.

The implementation of Milo is based on Milopy (https://github.com/emdann/milopy). It uses the same MuData based data structure that the scCODA 2.0 and tascCODA 2.0 implementations also use. Here, neighborhood counts are stored in a slot in MuData for downstream usage.

#### Multicellular programs with DIALOGUE

Multicellular programs, or gene programs, refer to the complex regulatory networks and signal transduction pathways that govern the behavior, differentiation, and communication of cells. DIALOGUE^37^ is a matrix factorization method for identifying these specific gene expression patterns. The implementation of DIALOGUE in pertpy resembles the original implementation^37^. The main differences are

- The R implementation of MultiCCA has been replaced with a Python implementation of the original mathematical formulation^77^, found at https://github.com/theislab/sparsecca. In addition, the Python implementation also has the option to solve for the canonical covariate weights *`w`* using linear programming, allowing for concurrent instead of iterative optimization over the pairwise factor matrices. This results in weights which are consistent regardless of the order in which cell types are passed, which was not previously true.
- A novel gene identification method, referred to as extrema MCP genes, which selects cells at the extreme values of the MCP (cells with the top 10% and bottom 10% MCP scores in each cell type), then runs the *rank_genes_groups* function from scanpy with default parameters to perform a t-test between the two groups of cells to identify differentially expressed genes to provide adjusted p-values based on the number of tested genes.

#### Enrichment with BlitzGSEA

Gene set enrichment analysis determines whether predefined sets of genes, often associated with specific biological functions or pathways, show statistically significant, concordant differences in expression across two biological states or phenotypes. It is used to identify biological processes that are overrepresented in a ranked list of genes, typically arising from high-throughput experiments. This approach shifts the analysis focus from individual genes to the collective behavior of genes within predefined, functionally related groups, facilitating a deeper understanding of the biological mechanisms underlying observed changes. Pertpy provides access to a variety of metadata databases that provide gene sets whose enrichment can be tested for.

We generally followed the enrichment pipeline described in drug2cell^78^ to test for the enrichment of gene sets. This pipeline entails:

1. Fetching gene sets from databases.
2. Scoring gene sets by computing the mean expression of each gene group per cell.
3. Performing a differential expression test to get ranked gene groups that are up-regulated in particular clusters.
4. Determining enriched genes using a hypergeometric test on the gene set scores or using BlitzGSEA^69^.

#### Distances, metrics, and permutation tests

Distance metrics serve as an important baseline in two primary tasks in single-cell perturbation analysis: 1) identifying relative heterogeneity and response and 2) evaluating and training single-cell perturbation models. To this end, various commonly used distance metrics have been implemented to be easily applied to single-cell AnnData objects with accompanying perturbation or disease labels. In the following, we present the 16 distances, in order of performance according to Ji et al.^54^, that are implemented in pertpy. We use *x^k^* to denote the gene expression in cell *k*, and *x_i_* and *y_i_* for the expression of gene *i* in the perturbed and control conditions, respectively.

**● Mean squared error (MSE)**

Determines the mean squared distance between the mean vectors of two groups.

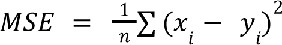

• **Maximum mean discrepancy (MMD)**

Evaluates the discrepancy between the empirical distributions of two groups using kernel-based methods. Let *n* denote the number of samples, and *k*(·, ·) the linear kernel function.

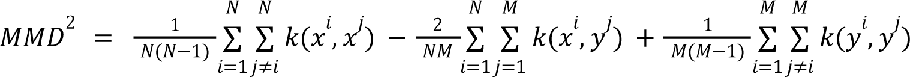

• **Euclidean distance**

Calculates the Euclidean distance between the means of the two groups.

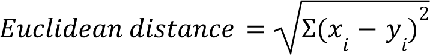

• **Energy distance**^39^

Computes a statistical energy distance between two groups based on mean pairwise distances within and between groups.

We define

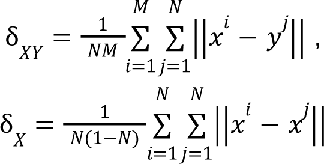

and δ*_Y_* accordingly. The energy distance is then calculated as

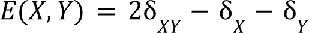

• **Kolmogorov-Smirnov Test distance (KS Test)**

Applies the KS statistic to measure the maximum distance between the empirical cumulative distributions of two groups. We define the empirical distribution function for gene *i* as

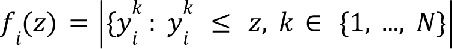

over all cells of the control condition and analogously 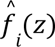 for perturbed cells. For each gene, the maximum distance between both distribution functions 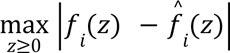 is computed, and the results averaged over all genes to yield a single distance value.

• **Mean absolute error (MAE)**

Measures the mean absolute difference between the mean vectors of two groups.

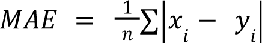

• **Two-sided T-test statistic**

Uses the T-test statistic to compare the means of two groups under the assumption of unequal variances. Let 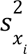 and 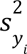 denote the variances of gene *i* for perturbed and control, *n_x_* and *n_y_* the sample sizes for perturbed and control, and ɛ a small factor to avoid dividing by zero.

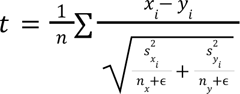

• **Cosine distance**

Computes the cosine of the angle between the mean vectors of the two groups.

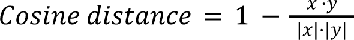

Where · denotes the dot product.

• **Pearson’s distance**

Uses Pearson correlation to assess the linear correlation between the mean vectors of two groups, returning 1 minus the correlation coefficient. Let 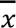 and 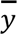 denote the mean expression over all genes.

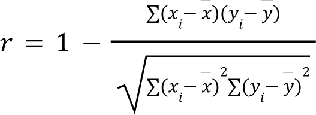

• **Coefficient of determination (R²) distance**

Calculates the coefficient of determination between the mean vectors of two groups. Note that, unlike most other distances listed here, R^2^ is not symmetric/has not been symmetrized.

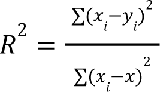

Where 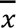 is the mean expression over all genes in the perturbed condition.

• **Classifier control probability**

To compute the classifier class projection distance between perturbations *P* and control condition *C*, we train a linear regression classifier to distinguish between *C* and *P*, with 20% of *P* held out for testing. To calculate the distance for perturbation class *P*_*i*_, we obtain the average post-softmax classification probabilities of all cells in *P*_*i*_ and return the probability of class *C*.

• **Kendall tau distance**

Applies Kendall’s tau, a measure of ordinal association, between the mean vectors of two groups. We define *C* as the number of concordant pairs, *D* as the number of discordant pairs, *X* as the number of ties in *x*’s ranking, and *Y* as the number of ties in *y*’s ranking.

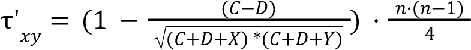

• **Spearman’s rank distance**

Similar to Pearson’s distance, but uses Spearman rank correlation to measure non-linear relationships.

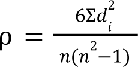

Where *d_i_* represents the difference in rank of gene *i* across both samples.

• **Wasserstein distance**

Also known as Earth Mover’s Distance, computes the cost of optimally transporting mass from one distribution to another. Let *W*(*p*, *q*) be the first order Wasserstein distance between probability distributions *p* and *q*, Γ(*p*, *q*) the set of all joint distributions with marginals *p* and *q*, *c*(*x*, *y*) the cost of transporting a unit of mass from *x* to *y*, and *X* and *Y* are the support sets of *p* and *q*, respectively.

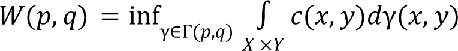

• **Symmetric Kullback-Leibler (KL) divergence**

Measures how one probability distribution diverges from a second. In the case of discrete inputs, the KL divergence is calculated as follows:

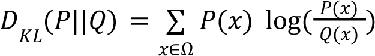

Where *P* and *Q* are discrete probability distributions.

For non-discrete inputs, the KL divergence is computed as

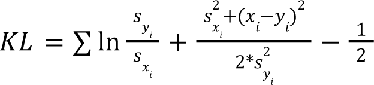

Where *s* denotes the standard deviation.

• **Classifier class projection**

The classifier class projection distance between perturbation *P_i_* and control condition *C_i_* is calculated by training a linear regression classifier on all *x* ∉ *P_i_* and all *C*, subsequently retrieving the average post-softmax classification probabilities of all cells 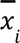, and returning the probability of class *C_i_*.

The following distance was also implemented in pertpy but were not part of the aforementioned benchmark:

• **Negative Binomial Log Likelihood (NBLL)**

Fits a negative binomial distribution to one group and uses it to compute the log likelihood of the other group’s data. For each gene *i* which is not overdispersed in *x*, we fit a negative binomial distribution with parameters µ_*i*_ and θ_*i*_. The distance between two categories *x* and *y* is then computed as the average negative log likelihood of *y* given the parameters of the distribution fit on *x* for each gene *i*, that is,

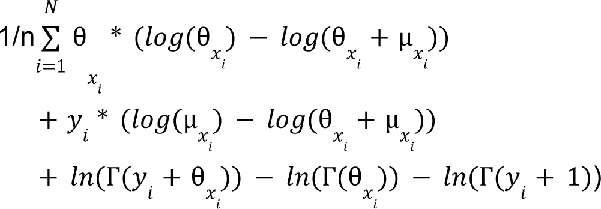

The *distances* module allows users to quickly fetch the pairwise distances between any set of categorically labeled cells. The *distance_tests* module allows users to compute a p-value through Monte-Carlo permutation testing, thereby providing a confidence value for any given distance. This can be particularly comforting in cases in which distances have been used as proxies for real biological response in gene expression space.

Note that while we refer to all of the above as “distances,” they do not all meet the mathematical definition of a distance; deviations from the standard distance axioms have been detailed in Ji et al^54^. While these distances can be used with any single-cell measurement, it should be noted that the ranking above was performed in the context of single-cell transcriptomics.

We also implemented two metrics for evaluating expression prediction models. To evaluate if perturbation prediction leads to meaningful biological conclusions we implemented a differential expression correlation metric. This metric uses Spearman correlation to compare differential gene ranking from the scanpy *rank_genes_groups* function performed on control vs real perturbed data and control vs predicted perturbed data. To evaluate if the distribution of gene expression means vs variances corresponds to real data we used a similar method as proposed before^79^. The distribution of expression mean-variance 2D relationship was estimated with kernel density for both real and predicted perturbed data. The distance between the two densities was estimated based on the difference of values sampled across the whole data range.

#### Perturbation ranking with Augur

Augur aims to rank or prioritize cell types according to their response to experimental perturbations. The fundamental idea is that in the space of molecular measurements cells reacting heavily to induced perturbations are more easily separated into perturbed and unperturbed than cell types with little or no response. This separability is quantified by measuring how well experimental labels (e.g. treatment and control) can be predicted within each cell type. Augur trains a machine learning model predicting experimental labels for each cell type in multiple cross validation runs and then prioritizes cell type response according to metric scores measuring the accuracy of the model. For categorical data Augur uses the area under the curve and for numerical data it uses concordance correlation coefficient.

Our implementation of Augur follows the original implementation^80,81^. We further optimized it by parallelizing the training of the predictive models. Moreover, the pertpy implementation allows for gene selection using either the originally used variance based implementation or scanpy’s highly variable genes.

#### Causal identification of single-cell experimental perturbation effects with CINEMA-OT

Cellular responses to environmental signals are crucial for understanding biological processes. Effectively extracting biological insights from such data, especially through single-cell perturbation analysis, remains challenging due to a lack of methods that can directly account for underlying confounding variations. CINEMA-OT distinguishes between confounding variations and the effects of perturbations, achieving an optimal transport match that mirrors counterfactual cell pairings. These pairings allow for the analysis of causal perturbation responses, enabling novel approaches including individual treatment-effect analysis, clustering of responses, attribution analysis, and the examination of synergistic effects.

The implementation of CINEMA-OT is based on the original implementation^38^. To further accelerate and simplify the implementation we used ott-jax^68^.

#### Perturbation spaces

Pertpy discriminates between two fundamental domains to embed and analyze data: the “cell space” and the “perturbation space”. In this paradigm, the cell space represents configurations where discrete data points represent individual cells. Conversely, the perturbation space departs from the individualistic perspective of cells and instead categorizes cells based on similar response to perturbation or expressed phenotype where discrete data points represent individual perturbations. This specialized space enables comprehending the collective impact of perturbations on cells. We differentiate between perturbation spaces (where we create one data point for all cells of one perturbation) and cluster spaces (where we cluster all cells and then test how well the clustering overlaps with the perturbations).

##### Pseudobulk Space

This space takes the pseudobulk of a covariate such as the condition to represent the respective perturbations using the Python implementation of DecoupleR^82^ (https://github.com/saezlab/decoupler-py) which can subsequently be embedded.

##### Centroid Space

The Centroid Space calculates the centroids as the mean of the points of a condition for a pre-calculated embedding. Next, it finds the closest actual point to that centroid which determines the perturbation space point for that specific condition.

##### Multilayer Perceptron Classifier Space

The Multilayer Perceptron (MLP) Classifier Space trains a feed-forward neural network to predict which perturbation has been applied to a given cell. By default, a neural network with one hidden layer of 512 neurons and batch-normalization is created and trained using a batch size of 256. However, all these hyperparameters can be customized by the user to suit the specific requirements of the dataset. We account for class imbalances by oversampling perturbations with fewer instances. The MLP is trained using cross entropy loss until detection of overfitting (early stopping), or until it reaches the maximum number of epochs to train, set to 40 by default. To obtain perturbation-informed embeddings of the cells, the cell representations in the last hidden layer are extracted. Another perturbation space such as pseudobulk can be applied downstream to obtain a per-perturbation embedding if required. For creation and training of the MLP, we leverage the PyTorch library.

##### Logistic Regression Classifier Space

The Logistic Regression Classifier Space generates perturbation embeddings, as opposed to per-cell embeddings computed by the MLP classifier space. A logistic regression classifier is trained for each perturbation individually to determine if the respective perturbation was applied to a cell or not. Depending on user preference, the classifier can be trained on the high-dimensional feature space or on a precomputed embedding, such as one obtained through PCA. For each perturbation, we extract the coefficients of the logistic regression classifier, trained until convergence or reaching the maximum number of iterations (1000 by default), to derive a per-perturbation embedding. We use scikit-learn’s implementation for the logistic regression classifier.

##### DBScan Space

DBSCAN^83^ (Density-Based Spatial Clustering of Applications with Noise) is a clustering algorithm that identifies clusters in a dataset based on the density of data points, grouping together points that are closely packed while marking points in low-density regions as outliers. Pertpy’s implementation of a DBScan Space is based on scikit-learn’s DBScan implementation.

##### K-Means Space

K-means is a clustering algorithm that partitions a dataset into K distinct, non-overlapping clusters by minimizing the distance between data points and the centroid of their assigned cluster. It iteratively adjusts the positions of centroids to reduce the total variance within clusters, making it suitable for identifying spherical-shaped clusters in feature space. Pertpy’s implementation of a K-Means Space uses k-means clustering as implemented in scikit-learn.

##### Label transfer

Label transfer in single-cell analysis involves using annotations of a datasets to predict the states of unannotated data points leveraging similarities in gene expression patterns or nearest neighbors. Pertpy’s label transfer function uses PyNNDescent to find the closest neighbors for all data points and then uses majority voting to label unlabeled data points.

#### Metadata support

Pertpy provides access to several databases that contain additional metadata for cell lines, mechanisms of actions, and drugs. On request, the database content gets cached locally and the respective information gets stored in the appropriate slots of the passed AnnData object.

##### Cell line

Pertpy provides access to DepMap (https://depmap.org/portal/, version 23Q4) and Genomics of Drug Sensitivity in Cancer (GDSC)^26^. The following information can be obtained:

- **Cell Line Identification**: Comprehensive details such as cell line names, aliases, DepMap IDs, and Cancer Cell Line Encyclopedia (CCLE)^84^ names.
- **Genetic Information**: Data on genetic aberrations prevalent in cancer cell lines, including mutations, copy number alterations (CNAs), fusion genes, and comprehensive gene expression profiles.
- **Dependency Scores**: Quantitative assessments of gene essentiality that showcases the impact of specific genes on the viability of cancer cell lines.
- **Drug Sensitivity**: Detailed measurements of how cancer cell lines respond to various drugs, with metrics such as IC50 values providing insights into the effectiveness and potential toxicity of therapeutic compounds.
- **Lineage and Type**: Information categorizing cell lines based on their tissue of origin and the type of cancer they represent.
- **Molecular Subtypes**: Classifications based on detailed genetic, epigenetic, and proteomic analyses, which help in understanding the heterogeneity within and across cancer types.
- **Phenotypic Data**: Observations on cell growth rates and morphological characteristics, which can correlate with genetic traits and drug responses.
- **Genomic Profiling**: Includes high-resolution data from whole-exome and whole-genome sequencing efforts, offering a comprehensive view of the genetic landscape of cell lines.
- **Proteomics Profiling**: Protein intensity values acquired using data-independent acquisition mass spectrometry (DIA-MS) from DepMap Sanger.

##### Mechanism of Action

Pertpy provides access to The Connectivity Map (CMAP)^27^, also commonly referred to as CLUE (CMap and LINCS Unified Environment) which hosts the infrastructure. CMAP is a resource designed to help researchers discover functional connections between diseases, genetic perturbation, and drug action. The following information can be obtained:

- **Compound names**: The name of the compound of genetic perturbagen.
- **Mechanism of Action**: The specific biochemical interactions through which compounds exert their effects on cellular functions. This includes detailed descriptions of whether a compound acts as an inhibitor, activator, or modulator of particular molecular targets.
- **Target**: The sets of genes or proteins that directly interacted with or were affected by the perturbagen.

##### Drug

Pertpy provides access to pubchem^28^ using pubchempy (https://github.com/mcs07/PubChemPy). PubChem is a comprehensive resource for chemical information, primarily known for its vast database of chemical molecules. The following information can be obtained:

- **Chemical Identifiers:** Each chemical in PubChem is assigned unique identifiers, including CAS numbers, InChI strings, and SMILES notation.

Pertpy further provides access to the CHEMbl^29^ database. ChEMBL is a comprehensive database maintained by the European Bioinformatics Institute (EBI), part of the European Molecular Biology Laboratory (EMBL). It provides a vast collection of data on bioactive molecules with drug-like properties. The following information can be fetched:

- **Compounds**: The names of the compounds.
- **Targets**: The target gene sets of the compounds.

#### Benchmarking runtime

To evaluate computational efficiency, we measured execution time and resource consumption for three tools implemented in pertpy: Augur, mixscape, and scCODA 2.0. Following their respective tutorials, we developed corresponding scripts in their original implementation. These scripts were executed on a system equipped with 8 Intel(R) Xeon(R) Gold 6142 CPUs @ 2.60GHz and 768GB of RAM, with 16GB allocated specifically for these tasks, in a Linux environment. This setup ensured accurate and reproducible timing measurements. Each script was run three times to guarantee consistency. Timing was recorded excluding the import time, using Python’s “time” package and the native function ‘Sys.time()’ in R. The results were displayed in a box plot (**Supplementary Figure 1**), which compared the execution time in seconds across each tool and implementation.

### Use-cases

For the following analyses, we used the latest pertpy version 0.8.0. We deposited a full Conda environment to reproduce our results in the associated reproducibility repository together with all result tables of our analysis.

#### Analysis of the CRISPR screen dataset

We obtained the original dataset from the original publication together with the labels of the gene programs. The dataset contained 111255 cells and 19018 genes. We followed the standard scanpy preprocessing pipeline to log normalize the data, calculate 4000 highly variable genes, obtain PCA components, and embedded the data into a UMAP space for visualization purposes. Moreover, we scored cell cycle genes using the list of Tirosh 2016^85^.

Afterwards, we used pertpy’s implementation of mixscape to calculate a perturbation signature which we further embedded into UMAP space. Next, we applied mixscape to the perturbation signature to calculate the perturbation scores that are automatically binarized to assign successful and unsuccessful perturbations. We used scikit-learn to calculate the silhouette before and after applying mixscape. The Silhouette Score varies between −1 and +1, signifying that a higher score denotes good alignment of an object with its own cluster and a poor alignment with adjacent clusters.

We applied pertpy’s multilayer perceptron based Discriminiator Classifier to the corrected space and embedded the pseudobulk of the penultimate layer feature values with UMAP. Pertpy’s label transfer function was applied to the nearest neighbor graph. To identify gene programs affected by perturbations in an unannotated cluster in the UMAP, we performed gene set enrichment analysis on either up- or downregulated genes (adjusted p-value cutoff of 0.01) in the cluster of interest, identifying the top three up- and downregulated reactome^86^ pathways for the cluster.

#### Analysis of the chemical perturbation dataset

We obtained the dataset from the original publication of the study which already contained annotations of cell lines, cell line quality, channel, disease, dose units, dose values, and many more fields that are documented in our analysis notebook. We filtered out cells perturbed by CRISPR, leaving 154710 cells and 32738 genes of 172 cell lines, treated with 13 different drugs. We applied standard preprocessing by filtering genes that were present in less than 30 cells and log normalizing the counts. 4000 highly variable genes were computed using the *highly_variable_genes* function of scanpy and used as the basis for downstream analyses, except when examining viability-dependent and -independent drug responses.

Next, we fetched all available cell line metadata from Cancer Dependency Map (DepMap) and Genomics of Drug sensitivity in Cancer using pertpy to annotate the cell lines by their DepMap ID with cell lineages, compound targets, and mechanism of action using Connectivity Map (CMAP)^27^. We further added drug sensitivities of cell lines to anti-cancer therapeutics from Genomics of Drug Sensitivity in Cancer (GDSC)^26^.

Pseudobulks were generated using pertpy’s *PseudobulkSpace* function by perturbation. We used the expression of the cell lines labeled as “control” as base lines. Bulk RNA expression data was fetched from the Cancer Cell Line Encyclopedia using the data from the Broad institute via pertpy. We used pertpy to calculate row-wise correlations of the expression profiles of the cell lines to obtain Pearson correlation values and p-values.

Finally, we used pertpy to disentangle drug responses into components that are independent of and dependent on the sensitivity of a certain cell line to a drug. We followed the approach presented in the paper introducing the original dataset^16^, but replaced functionalities with pertpy’s own implementation whenever possible. While previous work focused on the drug trametinib, we here investigated treatment responses to dabrafenib. We used pertpy’s *annotate_from_gdsc* function to query the IC50 values for each cell line-drug combination using the GDSC2 database. Next, we computed the expression log-fold change (log-FC) between treated cells and control based on raw counts for each cell line individually, using pertpy’s implementation of EdgeR. Then, for each gene, the following linear regression model was fit:

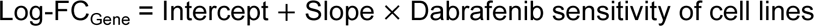

The fit model enables the decomposition of the observed change in gene expression in the treatment group into two components: a viability-independent response (intercept) and a viability-dependent response (slope). Genes with a Benjamini-Hochberg corrected p-value below 0.05 for either the slope or intercept were considered significant and subsequently used for gene set enrichment analyses using the gProfiler^87^ API.

#### Analysis of the TNBC treatment dataset

We obtained the dataset from the original publication which comprises scRNA- and ATAC-sequencing data from 22 patients with advanced triple-negative breast cancer (TNBC), treated with paclitaxel alone or in combination with the anti-PD-L1 therapy atezolizumab. We focused on the transcriptomic data which encompasses approximately 489,490 high-quality immune cells with 27085 measured genes across 99 high resolution cell types. We filtered genes with less than 10 cells, log normalized the data and selected highly variable genes using scanpy defaults. We calculated a PCA representation using scanpy with default settings that uses the “arpack” solver. For the following analyses, we filtered the dataset to only keep cell types that were retained in all response groups.

To determine compositional changes, we applied pertpy’s implementation of scCODA per treatment. scCODA’s automatic reference cell type detection determined intermediate monocytes as the reference cell type which we used for both treatments for consistency. Compositional changes with a false discovery rate of 0.1 (10%) were marked as credible effects.

We calculated the mean-squared error distance between the respective groups in a pairwise fashion using pertpy’s distance module on the PCA representation. We repeated this process three times for both treatments jointly, only chemotherapy treatment, and only anti-PDL1 and chemotherapy combination treatment. The results were visualized with seaborn.

DIALOGUE decomposition analysis was carried out exclusively on pre-treatment tumor samples. The sample labeled “Pre_P010_t” was excluded because it demonstrated low diversity in cell types. The analysis was confined to cell types that had a minimum of three cells per sample in the remaining patient samples. The number of MCPs was to set 10 with normalization enabled and the ‘LP’ solver.

A predictive MCP for treatment response was determined by individually testing each cell type within each MCP using a t-test for independent samples. To adjust for the number of cell types tested, the Benjamini-Hochberg correction method was applied. To identify significantly associated genes with the MCPs per cell type, cells at the extreme ends of the MCP distribution were selected, specifically, those in the top 10% and bottom 10% of MCP scores for each cell type. The scanpy *rank_genes_groups* function with default parameters was subsequently used. This function conducts a t-test between the two cell groups to pinpoint genes that are differentially expressed, offering an adjusted p-value that accounts for the total number of genes assessed. We filtered for heat shock proteins to determine HSPA1B to be significantly differentially expressed for Naive T cells (adj. P≤ 2.9e-272), CD8 effector memory cells (adj. P≤ 1.2e-172), CD4 regulatory T cells (adj. P≤ 5.3e-41), Plasma B cells (adj. P≤ 6.5e-34), CD4 central memory T cells (adj. P≤ 1.1e-01), and Memory B cells (adj. P≤ 6.5e-37)

To determine if the identified genes played a role in altered cell-cell interactions, gene comparisons were made for each cell type against the Nichenet database of protein-protein interactions, using gene names as identifiers^88^. An interaction was classified as MCP-associated if both the corresponding receptor and ligand were present among the significant genes (adjusted P-value less than 0.01) from two different cell types. An interaction was deemed MCP-ligand-associated if the ligand was linked to MCP in one cell type while the receptor exhibited a normalized mean expression over 1 in another cell type. Similarly, an interaction was considered MCP-receptor-associated if the receptor was connected to a MCP in one cell type and the ligand had at least 10 counts in the other cell type.

## Code and data availability

The pertpy source code is available at https://github.com/scverse/pertpy under the Apache 2.0 license. Further documentation, tutorials and examples are available at https://pertpy.readthedocs.io.

Scripts, notebooks, and analysis results to reproduce our analysis and figures are available at https://github.com/theislab/pertpy-reproducibility.

All used datasets are available through out-of-the-box dataloaders in pertpy.

## Acknowledgements

The authors thank all users of pertpy that regularly provide valuable feedback. We further thank the differential gene expression analysis team of the scverse hackathon in Cambridge in 2023 that developed prototypes and plots for the corresponding module in pertpy. We further thank Raphael Kfuri Rubens and Fabiola Curion for constructive comments on the manuscript.

## Author contributions

LH conceived the study. LH, YJ, XW, LM, AM, XZ, AS, JO, ED, MM, FC, IM, AM, SP, TG, AT, KH, MD, MB, IG, GS, AN, ER, ML, and SR implemented pertpy. LH and LM analyzed the CRISPR screen dataset. LM, LH, and YJ analyzed the chemical perturbation dataset. LM, LH, and TG analyzed the TNBC treatment dataset. ZZ and LH performed the runtime benchmarking. LH, LM, and YI wrote the manuscript. FJT, HBS, and CS supervised the work. All authors read, corrected and approved the final manuscript.

## Supplementary Material

### Supplementary Tables

**Supplementary Table 1.** DIALOGUE multicellular program adjusted p-values per cell type.

|  | MCP 1 | MCP 2 | MCP 3 | MCP 4 | MCP 5 | MCP 6 | MCP 7 | MCP 8 | MCP 9 | MCP 10 |
| --- | --- | --- | --- | --- | --- | --- | --- | --- | --- | --- |
| Memory B cells | 0.6829 | 0.1022 | 0.9515 | 0.9911 | 0.2550 | 0.7412 | 0.9577 | 0.8645 | 0.9990 | 0.7723 |
| CD4 central memory T cells | 0.4999 | 0.1022 | 0.9515 | 0.8197 | 0.3515 | 0.7412 | 0.9577 | 0.7628 | 0.9990 | 0.2804 |
| CD4 regulatory T cells | 0.4999 | 0.1022 | 0.9515 | 0.9761 | 0.2637 | 0.6213 | 0.9577 | 0.9589 | 0.9990 | 0.4192 |
| CD8 mucosal-associated invariant T cells | 0.4999 | 0.1392 | 0.9515 | 0.8674 | 0.2637 | 0.7880 | 0.9577 | 0.8214 | 0.9990 | 0.7723 |
| CD8 effector memory T cells | 0.4999 | 0.1115 | 0.9515 | 0.8674 | 0.6207 | 0.7880 | 0.9577 | 0.7628 | 0.9990 | 0.7723 |
| CD8 tissue-resident memory T cells | 0.4999 | 0.1392 | 0.9515 | 0.8197 | 0.6823 | 0.5370 | 0.9577 | 0.7628 | 0.9990 | 0.2804 |
| Naïve T cells | 0.4999 | 0.1022 | 0.9515 | 0.8197 | 0.2550 | 0.5370 | 0.9577 | 0.7628 | 0.9990 | 0.2804 |
| Intermediate monocytes | 0.9240 | 0.1115 | 0.9515 | 0.8197 | 0.6823 | 0.5370 | 0.9577 | 0.8962 | 0.9990 | 0.2804 |
| Plasma B cells | 0.9240 | 0.1022 | 0.9515 | 0.9911 | 0.2550 | 0.7880 | 0.9577 | 0.7628 | 0.9990 | 0.7723 |

### Supplementary Figures

**Supplementary Figure 1.**
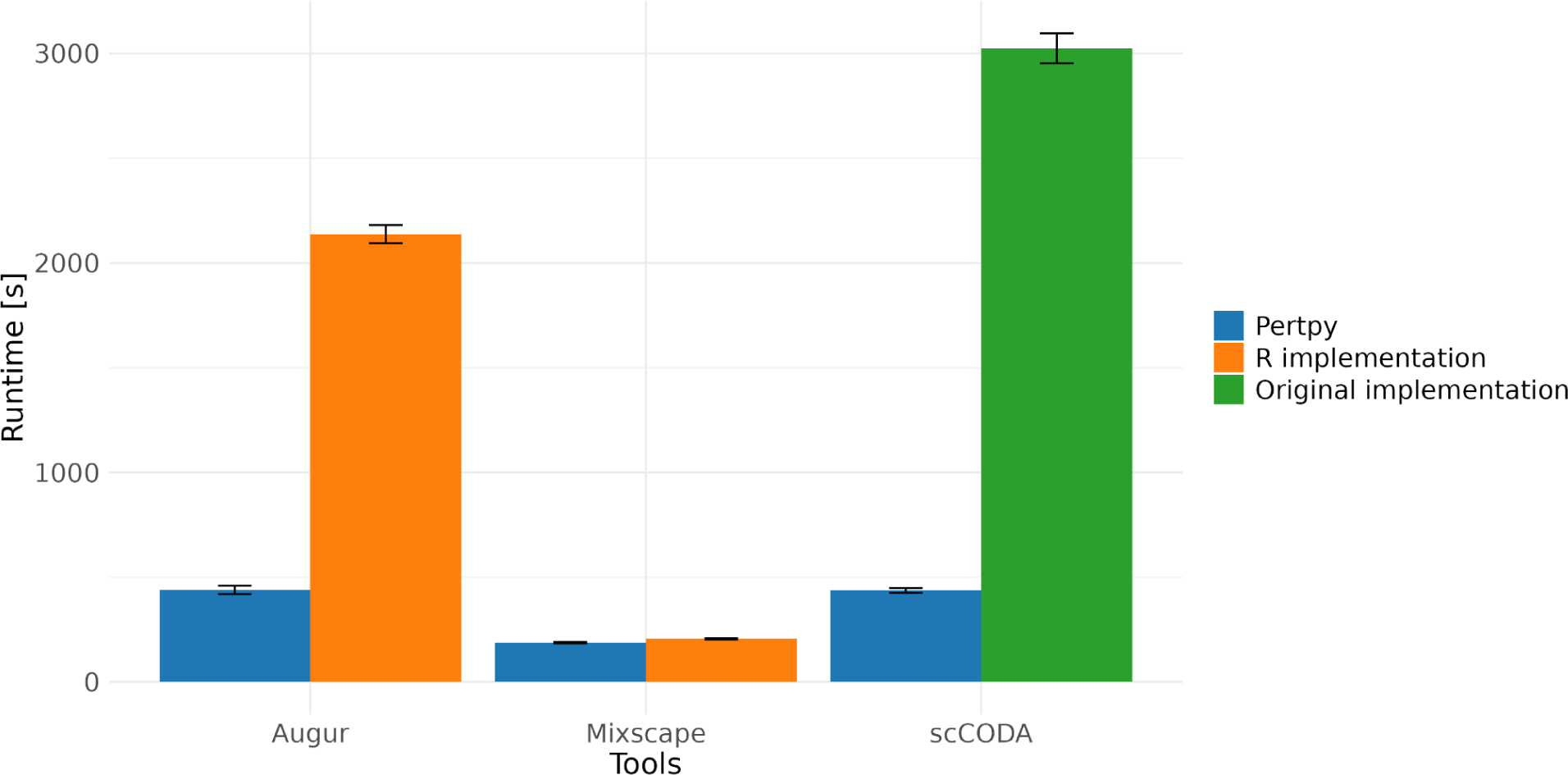
Runtime comparison of tools between pertpy’s implementation and correspondingly the existing R implementation or the formerly published original implementation. Tools were selected for benchmarking if the implementations differed substantially.

**Supplementary Figure 2.**
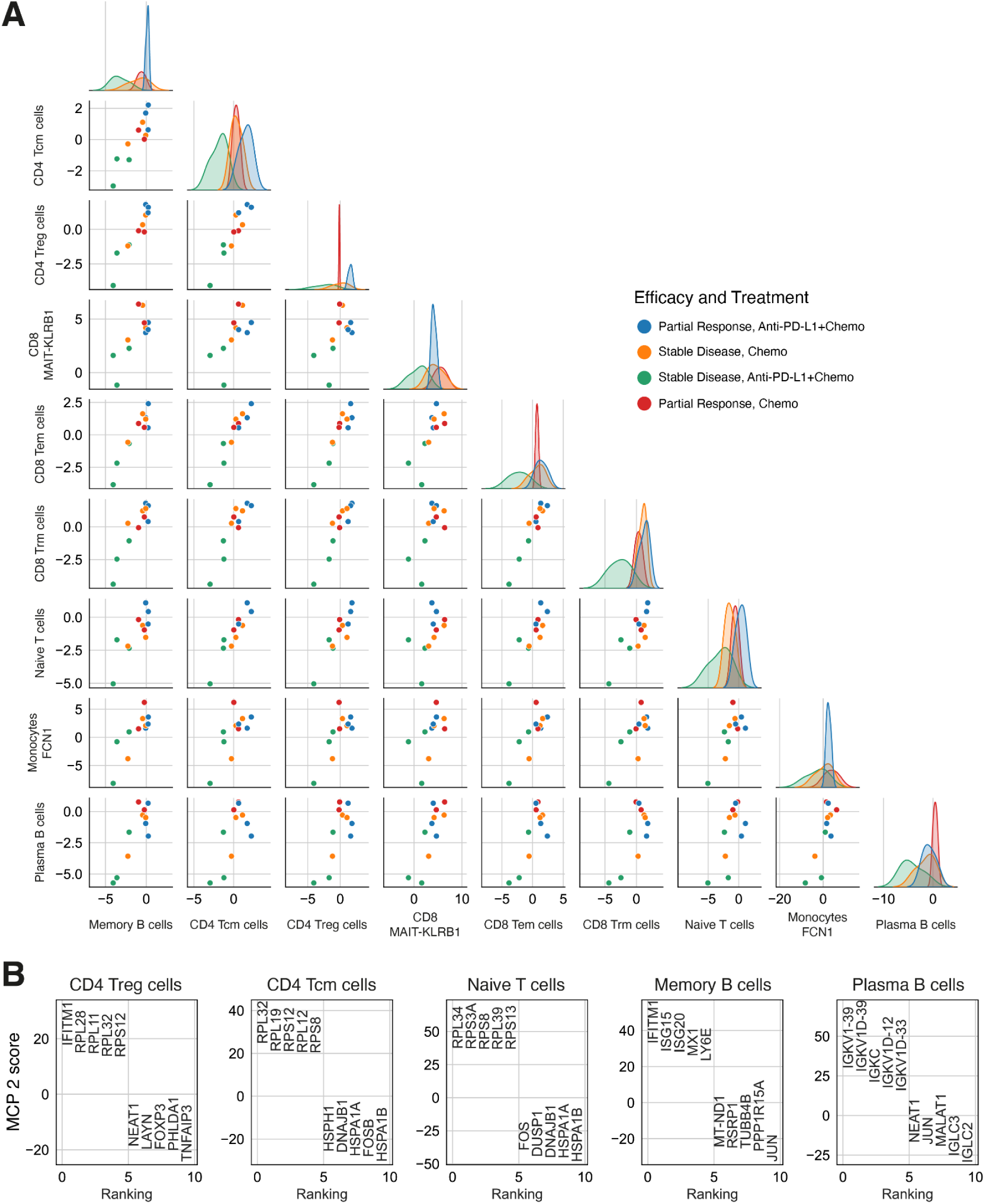
(A) Pair plots for MCP 2. The kernel density estimate along the diagonal shows the average score for each MCP by sample, specific to the indicated cell type. In the lower triangle’s scatter plots, each point signifies the average measurement from a patient for the cell types denoted by the respective row (x-axis) and column (y-axis). MCP 2 separates poor response to the PDL-1 inhibitors. (B) MCP 2 extrema genes per cell type. Shown are the respective five genes with the highest and lowest scores for MCP 2.

